# classLog: Logistic regression for the classification of genetic sequences

**DOI:** 10.1101/2022.08.15.503907

**Authors:** Michael A. Zeller, Zebulun W. Arendsee, Gavin J.D. Smith, Tavis K. Anderson

## Abstract

Sequencing and phylogenetic classification have become a common task in human and animal diagnostic laboratories. It is routine to sequence pathogens to identify genetic variations of diagnostic significance and to use these data in real-time genomic contact tracing and surveillance. Under this paradigm, unprecedented volumes of data are generated that require rapid analysis to provide meaningful inference. We present a machine learning logistic regression pipeline that can assign classifications to genetic sequence data. The pipeline implements an intuitive and customizable approach to developing a trained prediction model that runs in linear time complexity, generating accurate output more rapidly than other classification methods. Our approach was benchmarked against porcine respiratory and reproductive syndrome virus (PRRSv) and swine H1 influenza A (IAV) datasets. Trained classifiers were tested against sequences and simulated datasets that artificially degraded sequence quality at 0, 10, 20, 30, and 40%. When applied to a poor-quality sequence data, the classifier achieved between >85% to 95% accuracy for the PRRSv and the swine H1 IAV HA dataset and this increased to near perfect accuracy when using the full dataset. The model also identifies amino acid positions used to determine genetic clade identity through a feature selection ranking within the model. These positions can be mapped onto a maximum-likelihood phylogenetic tree, allowing for the inference of clade defining mutations. Our approach is implemented as a python package with code available at https://github.com/flu-crew/classLog.

## Introduction

Classification of pathogens has become a routine task in modern veterinary diagnostics (Shi et al. 2010). Classification of the infectious agent is a critical diagnostic step that allows for an informed decision on vaccination regimens and biosecurity measures that may be considered to clear a pathogen outbreak (Anderson et al. 2016; Kim et al. 2021; Paploski et al. 2019). Currently genetic classification is performed using phylogenetic methods such as maximum-likelihood and neighbor joining (Anderson et al. 2016; Chang et al. 2019; Turakhia et al. 2021). These methods are effective at classifying sequences and inferring relationships between taxa, but the time and skill required to execute and interpret analyses may impact their application in routine high-throughput activities. While diagnosticians are interested in the transmission and history of disease, the most pressing need is to provide a classification of data. Consequently, methods that do not conduct computationally intensive phylogenetic inference for inferring ancestry and genomic epidemiology are required.

Machine learning has been recognized as a viable method for classifying sequences (Kim et al. 2021; O’Toole et al. 2021). Genetic divergence over time leads to distinguishable genetic patterns within monophyletic clades that are linearly separable across aligned amino acid positions. This linear separability lends itself well to supervised machine learning methods such as logistic regression and random forest classification. Logistic regression based on aligned sequences is used as the primary means of automated classification for influenza A viruses (IAV) in swine that are processed within the FLUture database (Zeller et al. 2018). Similarly, porcine reproductive and respiratory syndrome virus (PRRSV) amino acid sequence data have been classified to genetic lineage using random forest, k-nearest neighbor, support vector machines, and multilayer perceptron methods (Kim et al. 2021). Decision tree machine learning approaches have been introduced to classify avian IAV sequences and SARS-CoV-2 sequences successfully at multiple taxonomical levels (Humayun et al. 2021; Randhawa et al. 2020). PangoLEARN, a random forest model, currently supplements the pangolin classification system for SARS-CoV-2 (O’Toole et al. 2021). However, despite machine learning appearing to be an effective approach for classification, few of these algorithms are user-friendly with intuitive generalized software that has been publicly released.

This manuscript introduces a general-purpose software application, classLog, that can train sequence classifiers based on user-labelled training data for use in classification of unknown sequences. The method used by the program leverages logistic regression, a parametric method of classification that runs in linear time complexity. Application of classLog provides a routine and robust way to integrate classification into pipelines where speed is necessary and there is no interest in inferring historical context of the sequences. Through decoupling the classification step from the inference of the history of the virus, this manuscript presents a method of classification that is rapid, accurate, and suitable for high-throughput pipelines.

## Methods

### Curation of swine H1 IAV and PRRSv North America datasets

We generated two datasets to test the utility of our classification pipeline: porcine reproductive and respiratory syndrome virus (PRRSV) and influenza A virus (IAV) in swine. We restricted the swine IAV to H1 subtype hemagglutinin (HA) genes from the United States collected between 2015 to 2021: these data were curated and annotated by genetic clade by the Influenza Research Database (Anderson et al. 2016; Zhang et al. 2017). Sequences sampled between 2015-2019 were used as a training set (n=3510), while 2020 and later sequences were extracted as a test set (n=163) (Figure 1B). For PRRSV sequences, we extracted the curated ORF5 gene sequence data provided by (Paploski et al. 2019), and extracted and assigned the genetic clade for each sequence from the GenBank accession’s feature information. The dataset was further refined by removing all “Type 1” European sequences, sequences that were not the full coding region, i.e., not equal to 603 or 606 nucleotides in length, and the remaining sequences were translated. The final dataset of 3047 annotated sequences were randomly split into training and test sets, using 80% (n=2,483) and 20% (n=609) of the sequences respectively (Figure 1A).

**Figure 1.**
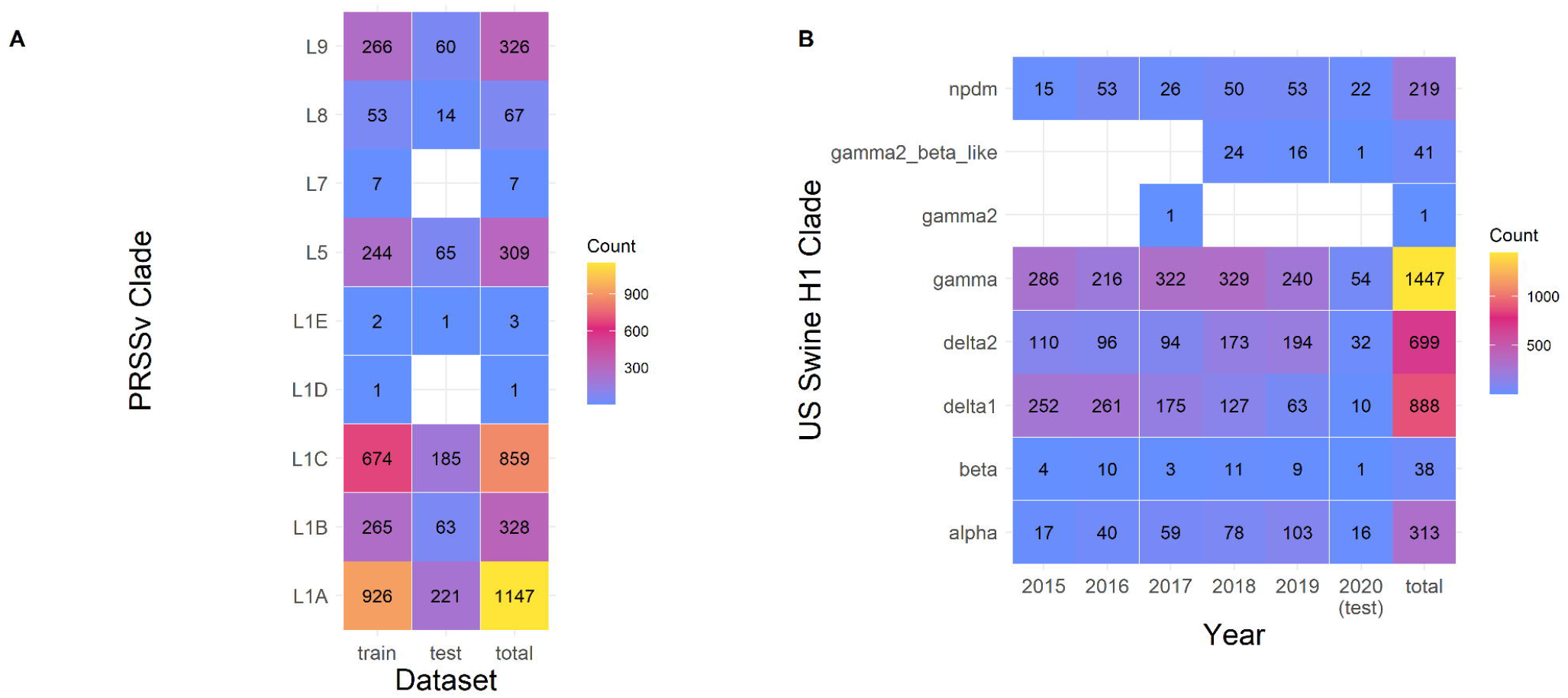
Number of samples in training and test sets for Porcine Respiratory and Reproductive Syndrome virus (PRRSv) and Swine H1 Influenza A virus (IAV). A) In the PRRSv dataset, number of samples were divided in a training set (80%) and a test set (20%). B) Samples in the IAV dataset per year. Samples collected in 2015-2019 were used as the training set, samples collected in 2020 were taken as the test set.

The datasets were split differently to simulate two distinct uses of the classifier. IAV data was split temporally to simulate classifying new data, while PRRSV sequences were split randomly to simulate filling in classifications from a mixed set.

### Simulated Sequencing Errors and Removing Informative Features

Gene sequences retrieved from Sanger sequencing, next generation, and third generation sequencing methods are not always complete, and there may be ambiguities and gaps in the data (Pfeiffer et al. 2018; Schirmer et al. 2016; Shendure and Ji 2008). These errors impact the estimation of the multiple sequence alignment that may subsequently decrease the accuracy of classification (Smirnov and Warnow 2021; Wang et al. 2009). To mimic decreasing quality of sequences, a python script was created to randomly generate a number of indices for replacement with an ambiguous ‘X’. Subsequently, the X’s were removed from the sequence to generate incomplete, unaligned sequences. Test set sequences were degraded at 0%, 10%, 20%, 30%, and 40% prior to classification. While more robust simulations of sequence degradation exist (Machado et al. 2021), the replication and implication of these methods is beyond the scope of this manuscript.

### Constructing classLog: the general sequence classifier

Sequence classification was implemented as a one-versus-rest logistic regression classifier. Input for classification requires an aligned nucleotide or amino acid FASTA file, with definition lines specifying the classification classes using character delimiters, e.g., A/swine/Iowa/A02636475/2022|1B.2.2.1, where ‘|’ delimits the phylogenetic clade from the strain name. The binary features of this model are the presence or absence of an amino acid at a specific position within the alignment. An optional feature selection process, which selects the most relevant sequence positions for classification, was implemented using a tree classifier that ranks binary features by GINI importance so that the user may restrict the prediction model to the most important features (Breiman et al. 1984). To facilitate the reusability of the classification scheme, the first sequence, feature labels, trained model, and class names are exported using a standard python pickle file format. The first sequence in the pickle file is used for pairwise alignment of unknown sequences to ensure there is consistency between query sequence alignment positions and the model feature positions. During prediction, a matrix of the presence or absence of nucleotides or amino acids at specific alignment positions is created, which is then fed to the model for prediction. For user submitted query sequences, the predicted classification is assigned and reported using classification names derived from the user-annotated classification fasta file used in training.

A prediction threshold option was included within the classifier to provide support for predicted classes on unknown data. Classifications with a score less than the threshold are rejected, and classified into an ‘unknown’ category (default value of 85%). The threshold criteria can have a direct effect on the performance of classification.

For validation, the general classifier was trained using 100%, 20%, 10%, 5%, 2%, 1%, and 0.5% of the available features within the H1 IAV and the PRRSv training datasets. For the H1 IAV sequence dataset, this resulted in 2439, 487, 243, 121, 24, and 12 features respectively. For the PRRSV dataset, this resulted in 686, 137, 68, 34, 6, and 3 features. Each classifier was used classify the 0%, 10%, 20%, 30%, and 40% test set sequences that had been generated to reflect sequencing errors and misalignment.

### Simplifying feature identification in query sequences using a Needleman-Wunsch pairwise alignment algorithm

An intrinsic challenge to the implementation of the machine learning classification process was correctly assigning the positions to new genetic sequences. To overcome this challenge without keeping the original alignment, a heuristic was applied such that the first sequence from the training set was saved and stored, and subsequent classification attempts would be pairwise aligned to recover the positions. To increase the speed and keep calculations within a tractable time for computation, a Needleman-Wunsch dynamic programming alignment algorithm (Needleman and Wunsch 1970) with affine gap penalties and a BLOSUM90 substitution matrix (Henikoff and Henikoff 1992) was implemented in C++ and exported as a python library using python bindings.

### Measuring the performance of the classifier on swine H1 IAV and PRRSv North America dataset classification

The performance of the classifiers was measured under the metrics of accuracy, macro precision, macro recall, and macro F1 (Jiao and Du 2016; Opitz and Burst 2019; Yang and Liu 1999). From a confusion matrix *M* where true classification is assigned along the y-axis and the predicted class is assigned along the x-axis, the precision and recall equations can be generalized as follows:

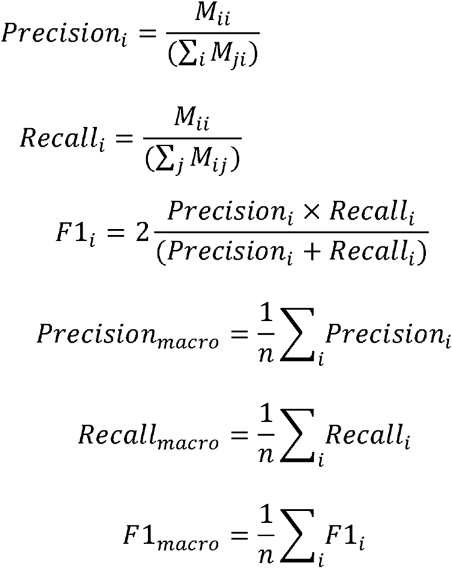

These metrics were taken for each classifier applied to the 0%, 10%, 20%, 30%, and 40% test set sequences with the results plotted using ggplot2 (Wickham 2016) in R v3.959 (R Core Team 2015).

### Visualization of swine H1 IAV and PRRSv North America dataset using ordination and phylogenetic analysis

Sequences from both datasets were aligned using MAFFT v7.487 (Katoh and Standley 2013). The pairwise number of differences between each sequence were extracted from the alignment using Geneious Prime 2022 (Kearse et al. 2012). These distances were ordinated into two-dimensional space using metric multidimensional scaling. Each ordination was colored first by the designated genetic clade, and then by a genetic motif consisting of the amino acids of the top two ranking amino acid positions. Amino acid position rank was calculated as the sum of GINI importance given by the extra tree classifier for each amino acid position, i.e., the two most important amino acids in determining the classification of the query sequence.

To identify the biological basis of the H1 swine IAV and PRRSv classifications, maximum likelihood trees were inferred for each dataset. Sequences were aligned using MAFFT v7.487 (Katoh and Standley 2013), and trees were inferred using IQ-TREE v1.6.12 (Nguyen et al. 2015). The PRRSv dataset was analyzed using a BLOSUM62 amino acid substitution model, while the IAV dataset was analyzed using the FLU amino acid substitution model (Dang et al. 2010). Statistical support was determined using the rapid bootstrap algorithm with 1,000 bootstraps, and the support was displayed on the branch of the resultant trees. Each tree was colored along the backbone by the phylogenetic clade, while the tips were annotated and colored by the top two ranking amino acid positions determined using GINI importance.

## Results

### classLog performance on H1 swine IAV and PRRSv observed and simulated data

A classLog classifier was trained on PRRSv ORF5 sequences collected and classified to lineage (Paploski et al. 2019), dividing the dataset into 80% training and 20% testing. The classifier performed perfectly correct when trained with 10% of features (n=68) of the total features with no sequence degradation (Figure 2A). At 10% sequence degradation (20aa), 10% of the features were able to achieve an accuracy of 97%. At 20% sequence degradation (40aa), 10% of the features were sufficient to achieve 88% accuracy, though increasing the number of features did not improve accuracy. Accuracy rapidly decreased at 30% sequence degradation (60aa), with 10% of the features achieving 69% correct classifications. At 40% sequence degradation (80aa) the greatest accuracy achieved was 42%.

**Figure 2.**
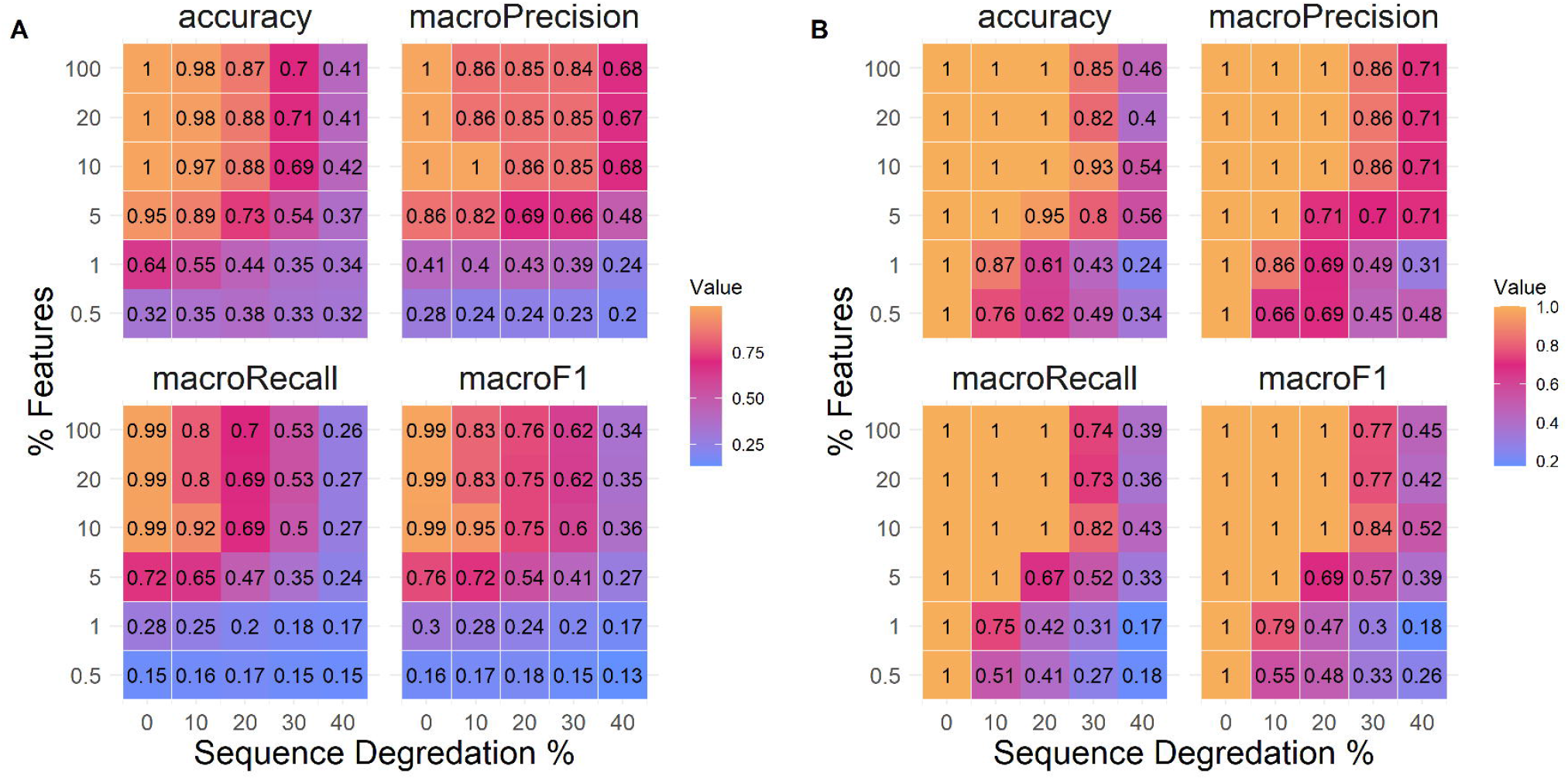
Measures of logistic regression classifier performance in the metrics of accuracy, precision, recall, and F1 scoring for the A) Porcine Respiratory and Reproductive Syndrome virus and B) Swine H1 Influenza A virus datasets. Each metric was measured over simulated sequence degradation of 0%, 10%, 20%, 30% and 40%, as well as with classifiers using 0.5%, 1%, 5%, 10%, 20%, and 100% of the available features for classification.

A classLog classifier was trained on H1 swine sequences present in IRD collected between 2015 to 2019, and was tested on 136 test sequences from 2020. The classifier performed perfectly correct when trained with as few as 12 features (0.5%) when there was no sequence degradation (Figure 2B). At 10% sequence degradation (56 aa), 5% of the features (121 features) were needed to achieve perfect accuracy. At 20% sequence degradation (112 aa), 10% of the features (243 features) were sufficient to achieve perfect accuracy. At 30% sequence degradation (170aa), 10% of the features were sufficient to achieve 93% correct classifications, although 20% of features (487 features) only achieved 82% correct classification. At 40% sequence degradation (227aa), there was a steep decline in the accuracy, falling below 60% across the board.

For both datasets, precision was consistently higher than recall (Figure 2). This is a consequence of rejecting classifications below the 85% scoring threshold and classifying them as ‘unknown,’ i.e., the number of false positives decreased while increasing false negatives.

### Using classLog to identify genetic features of biological relevance

The pairwise differences between the test set sequences were used to ordinate points in two-dimensional space (Figure 3). The ordination of both the PRRSv ORF5 and swine H1 IAV datasets were colored by their original designated clades, and by the motif formed by the amino acids present at the top two features ranked by GINI importance (Supplemental Figure 1 and 2). This manuscript uses the top two features as the number of amino acid combinations above two exceeds the number of distinct colors available on the pallet; but lower ranked features are important to discriminate between phylogenetic clades. Qualitatively, the ordination demonstrated separation between distant genetic lineages such as the H1 1A classical swine lineage versus the H1 1B human seasonal lineage (Figure 2C; Anderson et al 2016). However, sequences within some closely related genetic clades within the same lineage appeared to have overlap when assessed in a two-dimensional ordination. Within the PRRSv data (Figure 2B), the top two ranked amino acid positions (170, 172) corresponded well with the classified genetic clades suggesting that these positions may be clade defining mutations. For example, L1A has primarily the EE motif, L1B has EN, and L1C has DG. These divisions were not exclusive as L5, L8, L9 also have the EE motif that was exclusively within the L1A, and more features may need to be accounted for to discriminate between these clades. The top two positions of the swine H1 IAV dataset were 159 and 158 (H1 numbering, 17AA signal peptide removed) (Figure 2D), with a relatively high number of amino acid polymorphisms between those two positions. While some clades were well matched to one or two motifs, some clades such as alpha were highly varied in the motifs they carried, suggesting that other features position with a lower rank may better segregate this clade from the other clades. These data can be generated by extracting the features and their rankings using the classLog algorithm.

**Figure 3.**
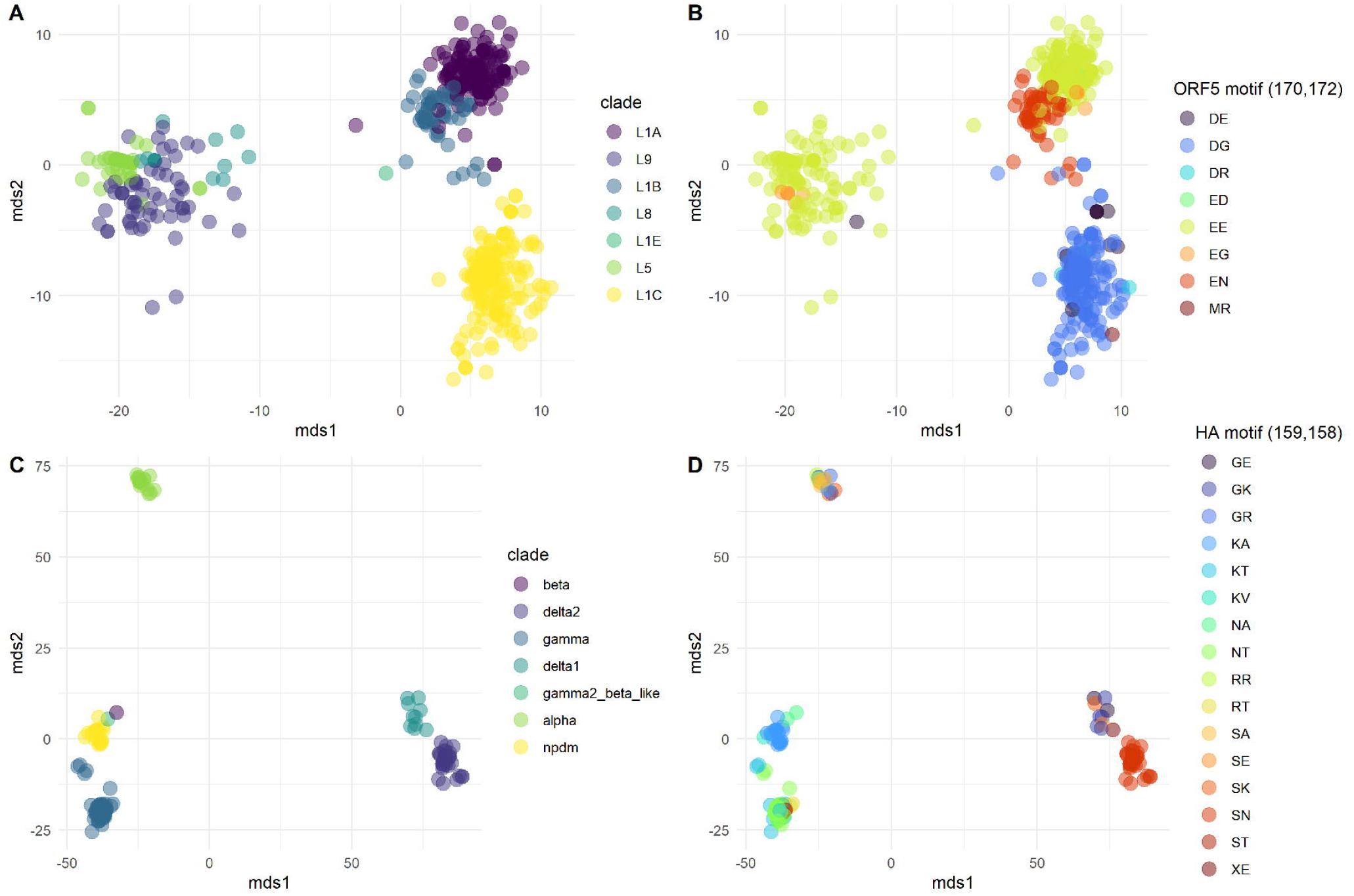
Metric multidimensional scaling in two dimensions of the number of pairwise differences between sequences of Porcine Respiratory and Reproductive Syndrome virus (PRRSv) ORF5 protein (A and B) and Swine H1 Influenza A virus datasets (IAV) (C and D). Plots were colored by genetic clade (A and C), and by the motif formed by the top two important positions inferred by decision tree (B and D). For PRRSv the top ranking features were positions 170 and 172. For IAV, the top ranking features were positions 159 and 158.

### Congruency between phylogenetic classification, classLog predictions, and model features

Maximum-likelihood trees were inferred for the PRRSv ORF5 and swine H1 IAV HA test datasets. The backbones of the phylogenetic trees were colored by the assigned genetic lineage, while the tips were labeled and colored by the motif formed by the two amino acid positions that had the highest cumulative GINI importance. For the PRRSv ORF5 dataset (Figure 4A: positions 170 and 172), the majority of L1B motifs were represented by an EN and L1C by DG. L1A, L5, L8, and L9 were also represented by EE at 170 and 172, suggesting that despite good concordance between the inferred phylogeny and the classLog predicted clade, this was being driven by features outside of these two positions.

**Figure 4.**
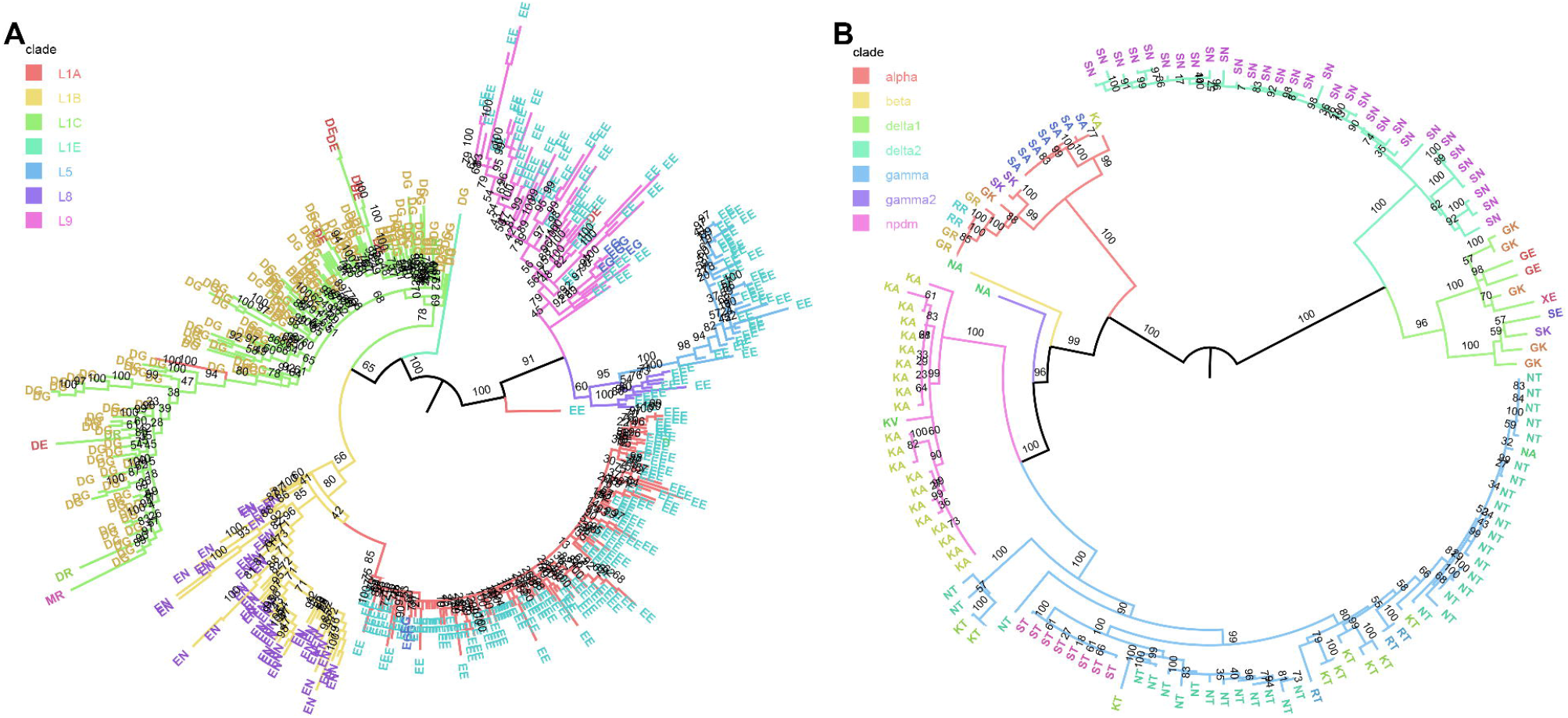
Phylogenetic maximum likelihood trees of A) Porcine Respiratory and Reproductive Syndrome virus (PRRSv) ORF 5 protein and B) Swine H1 Influenza A virus (IAV) test sequences, inferred by IQ-TREE v1.6.12 with 1,000 fast bootstraps. Tree backbones are colored by the prior assigned genetic lineage, where tip labels are colored by the motif formed by the top two ranking positions inferred by decision tree, positions 170 and 172 for PRRSv, and positions 159 and 158 for IAV. Bootstrap support is annotated on the branches.

For the swine H1 HA dataset (Figure 4B), the two most important features identified by classLog were positions 159 and 158. The majority of the 1B delta1a clade was primarily represented by GK, the 1B delta2 by SN, and 1A pandemic09 by KA. Three distinct motifs were identified within the 1A gamma clade, KT, NT, and ST, with RT interspersed. The 158T at was distinct enough to serve as a general rule to separate diversity within the 1A gamma clade. The remaining major H1 clade, 1A alpha, was associated with a significant amount of motif diversity, exhibiting GK, GR, KA, SA, SK and RR. The high amount of motif diversity is suggestive that another set of features may be used by the classifier for identifying this clade.

## Discussion

Applications of machine learning present computationally efficient ways of classifying genetic sequences without relying on traditional phylogenetic methods. The direct utility of machine learning methods is in high-throughput diagnostic processes, where the primary objective is to assign classification and there is not an immediate interest in inferring the evolutionary history of the sequence in question. By decoupling the classification process from phylogenetic method, complexity and computational time are reduced. Machine learning methods have the additional benefit of being highly portable and reproducible with minimal effort once an initial prediction model is trained. Our command line interface, classLog, represents a user-friendly and validated tool that can ingest annotated genetic sequences, train a classification model, and generate predictions and associated confidence scores without extensive computational and machine learning training.

classLog can be applied for rapid classification of genomic data either on site or in field settings. The advent of rapid and portable sequencing such as minION Nanopore technologies has resulted in the generation of thousands of sequences with a critical need to identify what they are, and whether the sample represents an “unknown.” The classLog program can be easily adopted as part of a light-weight pipeline that can be used to do classification on the fly in the field (e.g., Rambo-Martin et al. 2020). The execution of classLog does not require significant computational resources, and our testing was conducted on regular Windows and MacOS laptops. Consequently, it can easily be integrated within mobile diagnostic stations that are functional within remote locations that may have minimal access to extensive computational resources or trained personnel (e.g., Hoenen et al. 2016; Quick et al. 2016).

A consequence of field genomic epidemiology and the integration of Nanopore technology has been an increase in sequence error rate relative to traditional Sanger sequencing (Laver et al. 2015; Delahaye and Nicolas 2021). Our testing with classLog on simulated datasets, where we introduced sequence errors, suggested that the inaccuracies do not dramatically reduce the accuracy of classification using this machine learning method. It was noted that the classification failure within the H1 sequence dataset occurred proportionally to the number of samples present in the training dataset. As the sequence errors increased, misclassifications began to occur first in the sequences that had the least clade representation in the training set. It is likely that if there are more samples present in the training data to represent a specific clade, then the prediction model was better able to generalize the clade. This indicates one potential drawback of classLog, and that user-curated training datasets must remain large enough for optimal classifier performance. An alternate approach to generating large, curated datasets could be the application of classification models beyond the classLog logistic regression. Logistic regression is a parametric model that performs well on linearly separable classifications. In cases where the data are not linearly separable and that have limited training data, non-parametric models like SVM with an RBF kernel or random forest may perform significantly better, potentially provide easy to understand biological context to feature rankings (Sun et al. 2013; Zeller et al. 2021), but require more computational time and effort.

classLog is a method of creating light weight classifiers that can assign taxonomic classifications rapidly with minimal user curation and training. The implementation of this classification methodology can benefit diagnostic labs by saving computational run time associated with current phylogenetic classification approaches, and can be easily customized to work for different pathogens. An additional benefit is the identification of critical genetic features associated with clade classifications: these features are likely clade defining mutations and can be used to form hypotheses to investigate the gene to phenotype link (Abente et al. 2016; Koel et al. 2013; Lewis et al. 2014) and other functional studies. A benefit of machine learning approaches is that the results are also more directly interpretable as they are given as an assignment, rather than needing to be inferred from a tree. The culmination of these benefits offers a more streamlined approach to taxonomic assignment in a diagnostic setting.

## Acknowledgements

We gratefully acknowledge pork producers, swine veterinarians, and laboratories for participating in the USDA Influenza A Virus in Swine Surveillance System and publicly sharing sequences.

## Funding

This work was supported in part by: the U.S. Department of Agriculture (USDA) Agricultural Research Service [ARS project number 5030-32000-231-000-D]; the National Institute of Allergy and Infectious Diseases, National Institutes of Health, Department of Health and Human Services [contract numbers 75N93021C00015 and 75N93021C00016]; the USDA Agricultural Research Service Research Participation Program of the Oak Ridge Institute for Science and Education (ORISE) through an interagency agreement between the U.S. Department of Energy (DOE) and USDA Agricultural Research Service [contract number DE-AC05-06OR23100]; the SCINet project of the USDA Agricultural Research Service [ARS project number 0500-00093-001-00-D]; and the Duke-NUS Signature Research Program funded by the Ministry of Health, Singapore. The funders had no role in study design, data collection and interpretation, or the decision to submit the work for publication. Mention of trade names or commercial products in this article is solely for the purpose of providing specific information and does not imply recommendation or endorsement by the USDA, DOE, or ORISE. USDA is an equal opportunity provider and employer.

### Supplementary data and data availability

The machine learning classifier and data associated with the analysis in this manuscript is provided at https://github.com/flu-crew/classLog. Supplementary figures are available online at XXX.

## Notes

https://github.com/flu-crew/classLog

### Competing Interest Statement

The authors have declared no competing interest.

## References

Abente, Eugenio J, et al. (2016), ‘The molecular determinants of antibody recognition and antigenic drift in the H3 hemagglutinin of swine influenza A virus’, Journal of virology, 90 (18), 8266–80.

Anderson, Tavis K, et al. (2016), ‘A Phylogeny-Based Global Nomenclature System and Automated Annotation Tool for H1 Hemagglutinin Genes from Swine Influenza A Viruses’, mSphere, 1 (6), e00275–16.

Breiman, Leo, et al. (1984), Classification and regression trees (CRC press).

Chang, Jennifer, et al. (2019), ‘octoFLU: Automated Classification for the Evolutionary Origin of Influenza A Virus Gene Sequences Detected in US Swine’, Microbiology resource announcements, 8 (32), e00673–19.

Dang, Cuong Cao, et al. (2010), ‘FLU, an amino acid substitution model for influenza proteins’, BMC evolutionary biology, 10 (1), 1–11.

Delahaye, C. and Nicolas, J. (2021), ‘Sequencing DNA with nanopores: Troubles and biases’, PLoS One, 16(10), p.e0257521.

Henikoff, Steven and Henikoff, Jorja G (1992), ‘Amino acid substitution matrices from protein blocks’, Proceedings of the National Academy of Sciences, 89 (22), 10915–19.

Hoenen T, Groseth A, Rosenke K, Fischer RJ, Hoenen A, Judson SD, Martellaro C, Falzarano D, Marzi A, Squires RB, Wollenberg K R. (2016), ‘Nanopore sequencing as a rapidly deployable Ebola outbreak tool’, Emerging Infectious Diseases, 22(2), 331.

Humayun, Fahad, et al. (2021), ‘Computational Method for Classification of Avian Influenza A Virus Using DNA Sequence Information and Physicochemical Properties’, Frontiers in Genetics, 12, 10.

Jiao, Yasen and Du, Pufeng (2016), ‘Performance measures in evaluating machine learning based bioinformatics predictors for classifications’, Quantitative Biology, 4 (4), 320–30.

Katoh, Kazutaka and Standley, Daron M (2013), ‘MAFFT multiple sequence alignment software version 7: improvements in performance and usability’, Molecular biology and evolution, 30 (4), 772–80.

Kearse, Matthew, et al. (2012), ‘Geneious Basic: an integrated and extendable desktop software platform for the organization and analysis of sequence data’, Bioinformatics, 28 (12), 1647–49.

Kim, Jeonghoon, et al. (2021), ‘Applications of Machine Learning for the Classification of Porcine Reproductive and Respiratory Syndrome Virus Sublineages Using Amino Acid Scores of ORF5 Gene’, Frontiers in Veterinary Science, 813.

Koel, Björn F, et al. (2013), ‘Substitutions near the receptor binding site determine major antigenic change during influenza virus evolution’, Science, 342 (6161), 976–79.

Laver, T., Harrison, J., O’neill, P.A., Moore, K., Farbos, A., Paszkiewicz, K. and Studholme, D.J. (2015), ‘Assessing the performance of the oxford nanopore technologies minion’, Biomolecular detection and quantification, 3, 1–8.

Lewis, Nicola S, et al. (2014), ‘Substitutions near the hemagglutinin receptor-binding site determine the antigenic evolution of influenza A H3N2 viruses in US swine’, Journal of virology, 88 (9), 4752–63.

Machado, Denis Jacob, Castroviejo-Fisher, Santiago, and Grant, Taran (2021), ‘Evidence of absence treated as absence of evidence: The effects of variation in the number and distribution of gaps treated as missing data on the results of standard maximum likelihood analysis’, Molecular phylogenetics and evolution, 154, 106966.

Needleman, Saul B and Wunsch, Christian D (1970), ‘A general method applicable to the search for similarities in the amino acid sequence of two proteins’, Journal of molecular biology, 48 (3), 443–53.

Nguyen, Lam-Tung, et al. (2015), ‘IQ-TREE: a fast and effective stochastic algorithm for estimating maximum-likelihood phylogenies’, Molecular biology and evolution, 32 (1), 268–74.

Opitz, Juri and Burst, Sebastian (2019), ‘Macro f1 and macro f1’, arXiv preprint 1911.03347.

O’Toole, Áine, et al. (2021), ‘Assignment of epidemiological lineages in an emerging pandemic using the pangolin tool’, Virus Evolution, 7 (2), veab064.

Paploski, Igor Adolfo Dexheimer, et al. (2019), ‘Temporal dynamics of co-circulating lineages of porcine reproductive and respiratory syndrome virus’, Frontiers in microbiology, 10, 2486.

Pfeiffer, Franziska, et al. (2018), ‘Systematic evaluation of error rates and causes in short samples in next-generation sequencing’, Scientific reports, 8 (1), 1–14.

Quick, J., Loman, N.J., Duraffour, S., Simpson, J.T., Severi, E., Cowley, L., Bore, J.A., Koundouno, R., Dudas, G., Mikhail, A. and Ouédraogo, N. (2016). ‘Real-time, portable genome sequencing for Ebola surveillance’, Nature, 530(7589), 228–232.

R Core Team (2015), ‘R: A language and environment for statistical computing’.

Rambo-Martin, Benjamin L, et al. (2020), ‘Influenza A virus field surveillance at a swine-human interface’, MSphere, 5 (1), e00822–19.

Randhawa, Gurjit S, et al. (2020), ‘Machine learning using intrinsic genomic signatures for rapid classification of novel pathogens: COVID-19 case study’, Plos one, 15 (4), e0232391.

Schirmer, Melanie, et al. (2016), ‘Illumina error profiles: resolving fine-scale variation in metagenomic sequencing data’, BMC bioinformatics, 17 (1), 1–15.

Shendure, Jay and Ji, Hanlee (2008), ‘Next-generation DNA sequencing’, Nature biotechnology, 26 (10), 1135–45.

Shi, Mang, et al. (2010), ‘Molecular epidemiology of PRRSV: a phylogenetic perspective’, Virus research, 154 (1-2), 7–17.

Smirnov, Vladimir and Warnow, Tandy (2021), ‘Phylogeny estimation given sequence length heterogeneity’, Systematic biology, 70 (2), 268–82.

Sun, Hailiang, et al. (2013), ‘Using sequence data to infer the antigenicity of influenza virus’, MBio, 4 (4), e00230–13.

Turakhia, Yatish, et al. (2021), ‘Ultrafast Sample placement on Existing tRees (UShER) enables real-time phylogenetics for the SARS-CoV-2 pandemic’, Nature Genetics, 53 (6), 809–16.

Wang, Li-San, et al. (2009), ‘The impact of multiple protein sequence alignment on phylogenetic estimation’, IEEE/ACM transactions on computational biology and bioinformatics, 8 (4), 1108–19.

Wickham, Hadley (2016), ggplot2: elegant graphics for data analysis (Springer).

Yang, Yiming and Liu, Xin (1999), ‘A re-examination of text categorization methods’, Proceedings of the 22nd annual international ACM SIGIR conference on Research and development in information retrieval, 42–49.

Zhang, et al. (2017), ‘Influenza Research Database: An integrated bioinformatics resource for influenza virus research’, Nucleic Acids Res, 45 (D1), D466–D74.

Zeller, Michael A, et al. (2018), ‘ISU FLU ture: a veterinary diagnostic laboratory web-based platform to monitor the temporal genetic patterns of Influenza A virus in swine’, BMC bioinformatics, 19 (1), 397.

Zeller, Michael A, et al. (2021), ‘Machine learning prediction and experimental validation of antigenic drift in H3 influenza A viruses in swine’, Msphere, 6 (2), e00920–20.

