## Supplemental Material for "classLog: Logistic regression for the classification of genetic sequences"

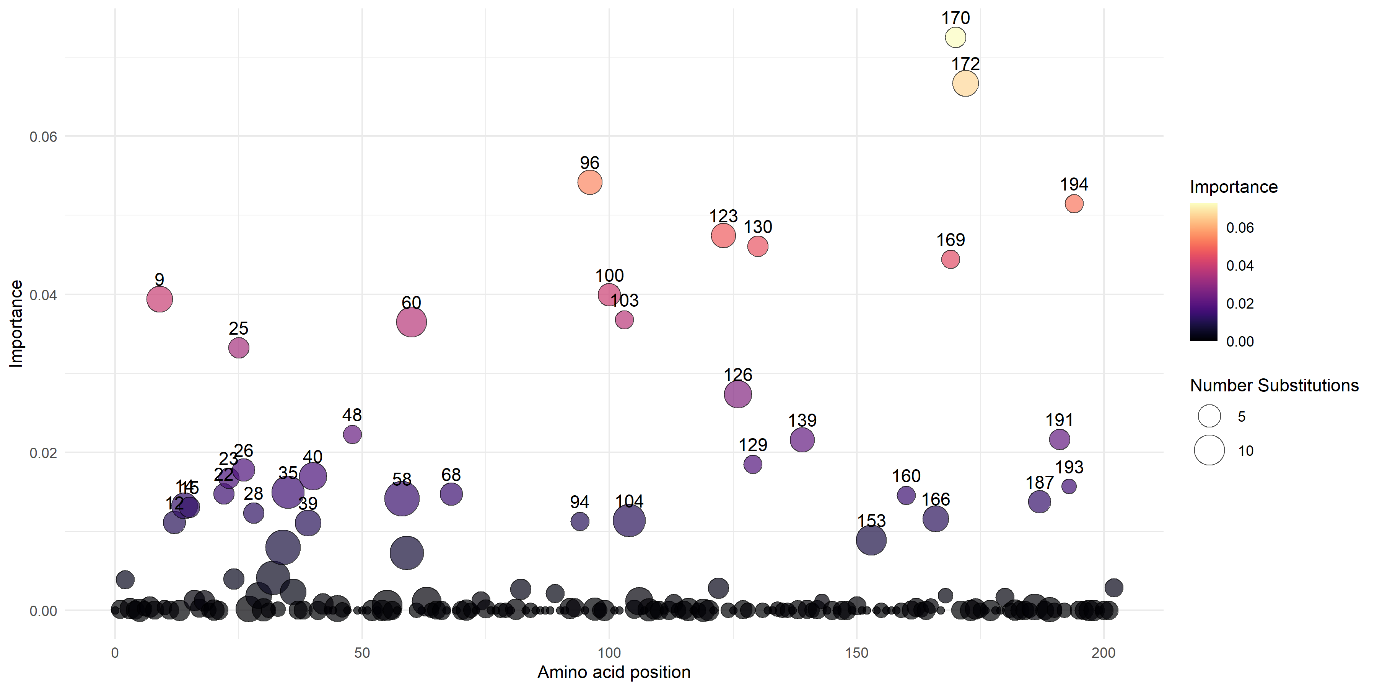

**Supplemental Figure 1** Cumulative summation of GINI importance at each amino acid position for the aligned Porcine Reproductive and Respiratory Syndrome virus (PRRSv) ORF5. The top 5 features in order are 170, 172, 96, 194, and 123. Color scales with importance, position size scales with the number of distinct substitutions observed at the position in the alignment.

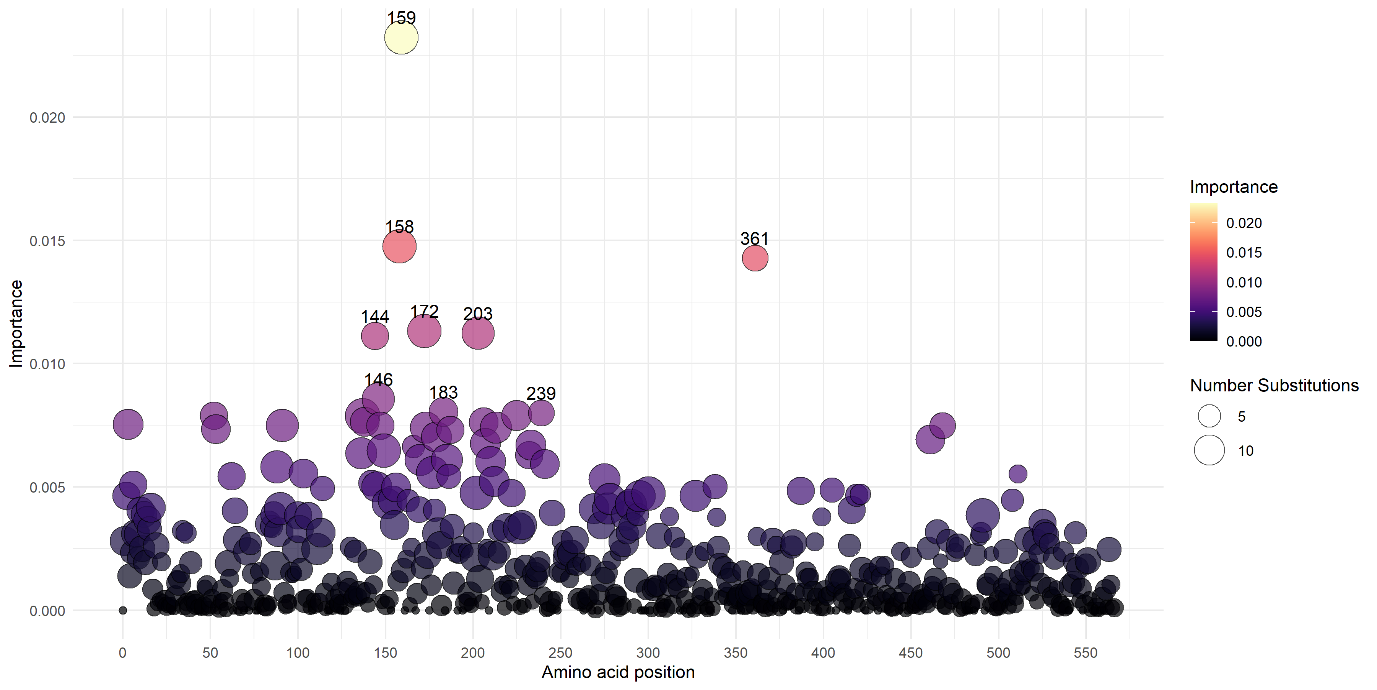

**Supplemental Figure 2** Cumulative summation of GINI importance at each amino acid position for the aligned Swine H1 Influenza A virus (IAV). The top 5 features in order are 159, 158, 361, 172, and 203. Color scales with importance, position size scales with the number of distinct substitutions observed at the position in the alignment.
